# First comprehensive identification of proteins with increased O-GlcNAc levels during pressure overload hypertrophy

**DOI:** 10.1101/2022.02.17.480962

**Authors:** Wei Zhong Zhu, Teresa Palazzo, Mowei Zhou, Dolena Ledee, Heather M Olson, Ljiljana Paša-Tolić, Aaron K. Olson

## Abstract

Protein posttranslational modifications (PTMs) by O-GlcNAc globally rise during pressure-overload hypertrophy (POH). However, only a few specific proteins with altered O-GlcNAc levels during POH are known primarily because this PTM is easily lost during standard mass spectrometry (MS) conditions used for protein identification. Methodologies have recently emerged to stabilize the O-GlcNAc moiety for MS analysis. Accordingly, our goal was to determine the proteins undergoing changes in O-GlcNAc levels during POH. We used C57/Bl6 mice subjected to Sham or transverse aortic constriction (TAC) to create POH. From the hearts, we stabilized and labelled the O-GlcNAc moiety with tetramethylrhodamine azide (TAMRA) before enriching by TAMRA immunoprecipitation (IP). We used LC-MS to identify the captured O-GlcNAcylated proteins. We identified a total of 707 O-GlcNAcylated proteins in Sham and POH. Two hundred thirty-three of these proteins were significantly increased in POH over Sham whereas no proteins were significantly decreased in POH. We examined two MS identified proteins, CPT1B and PDH, to validate the MS data by immunoprecipitation. We corroborated increased O-GlcNAc levels during POH for the metabolic enzymes CPT1B and PDH. Enzyme activity assays showed higher O-GlcNAcylation increased CPT1 activity and decreased PDH activity. In summary, we generated the first comprehensive list of proteins with changes in O-GlcNAc levels during POH and, to our knowledge, the largest list for any cardiac pathology. Our results demonstrate the large number of proteins and cellular processes affected by O-GlcNAc during POH and serve as a guide for testing specific O-GlcNAc-regulated mechanisms.

## BACKGROUND

Posttranslational modifications (PTM) of serine/threonine protein residues by O-linked β-N-acetylglucosamine (O-GlcNAc) are a dynamic and reversible process affecting protein functions [1, 2]. Diverging from most PTMs, protein O-GlcNAcylation is regulated by a single enzyme for attachment (O-GlcNAc transferase, OGT), and a single enzyme for removal (O-GlcNAcase, OGA). Thus, overall protein O-GlcNAc modifications increase or decrease in a more coordinated manner than most PTMs. Studies in non-cardiac tissues indicates protein O-GlcNAcylation modifies hundreds of proteins, but these results have not been confirmed in the heart [1].

Increased myocardial total protein O-GlcNAc levels during hypertrophy and heart failure were first noted about a decade ago [2]. Despite multiple studies confirming augmentation of overall protein O-GlcNAc levels during hypertrophy and heart failure in humans and animal models [3–8], only a few O-GlcNAc modified proteins or affected cellular processes have been identified [9, 10]. The lack of known O-GlcNAcylated proteins is partially due to the fragility of the O-GlcNAc moiety, which is lost during standard mass spectroscopy methodologies used to derive peptide sequence information and/or assign the modification site during proteomic studies [11].The O-GlcNAc moiety is also low abundance, so sample enrichment is required to detect these modified proteins [12]. These technical challenges have led to a critical knowledge deficit around the underlying effects from changes in O-GlcNAcylated proteins during cardiac hypertrophy.

Methodologies have recently emerged to stabilize the O-GlcNAc moiety on proteins and enrich O-GlcNAcylated proteins (peptides) allowing for their identification and quantitation using standard LC-MS techniques [12–14]. Accordingly, our goal was to create the first comprehensive list of proteins undergoing changes in O-GlcNAc levels during pressure overload hypertrophy (POH). We used our established model of POH from transverse aortic constriction (TAC) to increase total protein O-GlcNAc levels. From these hearts, we first stabilized and labelled the O-GlcNAc moiety with tetramethylrhodamine azide (TAMRA) [14]. We then enriched for O-GlcNAc modified proteins with a TAMRA immunoprecipitation and identified the captured proteins via LC-MS.

## MATERIALS AND METHODS

### Ethics statement

This investigation conforms to the Guide for the Care and Use of Laboratory Animals published by the National Institute of Health (NIH Pub. No. 85-23, revised 1996) and were reviewed and approved by the Office of Animal Care at Seattle Children’s Research Institute. All materials used in this study are available commercially from the indicated vendors.

### Mice

The studies were performed in male C57BL/6J mice from the Jackson Laboratory (Bar Harbor, ME) between the ages of 3 and 5 months. All mice were allowed free access to water and Teklad #7964 chow (Envigo, East Millstone, NJ, USA). Teklad ¼ inch corncob bedding was utilized in the cages (Envigo, East Millstone, NJ, USA).

### Experimental set-up

For the proteomics study, we compared hearts 1-week post-surgery from either TAC (POH) or Sham (Sham). We chose this duration after surgery because our prior study showed total protein O-GlcNAc levels were substantially increased by approximately 70% at this time [8].

To circumvent the issue of variable total protein levels observed between POH and Sham, we assessed enzymatic activity using the following model. Mice 4-weeks post-TAC were given intraperitoneal injections of the OGA inhibitor thiamet-g (TG) or vehicle (V) for 2 weeks. The groups for these experiments are TG-TAC and V-TAC and were previously detailed [8]. We showed total protein O-GlcNAc levels were significantly increased by about 4-fold in TG-TAC versus V-TAC [15].

### Surgery

We performed TAC or Sham surgery as previously described in our lab [7, 8, 16]. Briefly, mice were initially anesthetized with 3% isoflurane in 100% O2 at a flow of 1 LPM and then maintained with 1.5% isoflurane for the duration of the surgery. A midline sternotomy was performed to expose the aorta. The aorta was constricted with 7-0 silk suture tied against a 22-gauge blunt needle. Sham mice underwent similar surgery but did not have the suture tightened around the aorta. For pain control, mice received intraperitoneal buprenorphine (0.05-0.1 mg/kg IP) starting pre-operatively and continuing approximately every 12 hours for two and a half days.

### Echocardiograms

We performed echocardiograms as previously described in our lab [7, 16–18] to measure cardiac function and the degree of aortic constriction from the surgery. Briefly, mice were initially sedated with 3% isoflurane in O2 at a flow of 1 LPM and placed in a supine position at which time the isoflurane is reduced to 1.5% administered via a small nose cone. ECG leads were placed for simultaneous ECG monitoring during image acquisition. Mice were maintained a temperature of 37.5° C throughout the echocardiogram. Echocardiographic images were performed with a Vevo 2100 machine using a MS250 or MS550 transducers (VisualSonics, Inc, Toronto, Canada). M-Mode measurements at the midpapillary level of the left ventricle (LV) were performed at end-diastole (EDD) and end-systole (ESD) to determine LV function via the fractional shortening [(LVEDD-LVESD)/LVEDD * 100] in a parasternal short axis mode for at least three heart beats. We measured the pulse wave Doppler velocity across the aortic constriction site for a non-invasive estimate of the peak instantaneous pressure gradient across the constriction site as calculated by the simplified Bernoulli equation (Δ pressure gradient = 4*velocity^2^). The echocardiogram reader was blinded to treatment. We previously found that isoflurane anesthesia during echocardiograms increases myocardial total protein O-GlcNAc levels for several hours after the procedure (unpublished data), so the echocardiograms were always performed one day prior to sacrifice.

### Thiamet-G (TG)

We used the selective OGA inhibitor TG (Cayman Chemical Company, Ann Arbor, MI) at 20 mg/kg/day once daily for two weeks intraperitoneal to increase protein O-GlcNAc levels as previously shown [8]. TG was prepared in dimethyl sulfoxide (approximately 20 microliters) and the V-TAC received an equivalent volume of the vehicle only.

### Enzymatic labeling of O-GlcNAc-modified proteins for stabilization and enrichment

O-GlcNAc modified proteins were labeled using Click-iT™ O-GlcNAc Enzymatic Labeling System (Thermo Fisher Scientific, Waltham, MA) according to manufacturer’s instructions. Briefly, protein was extracted in RIPA lysis buffer plus phosphatase inhibitor (Thermo Fisher Scientific, Waltham, MA), the OGA inhibitors PUGNAc 20 µmol/L (Toronto Research Chemical) and Thiamet-G 1 µmol/L (Adooq Bioscience, Irvine CA). 200 µg of total protein was precipitated using chloroform/methanol to remove the detergents followed by centrifugation at 14,000g for 5 minutes at 4°C. The interface layer containing the protein precipitate was washed twice with methanol. The pellet was dried for 5 minutes in a standard fume hood. The dried pellet was resuspended in 40µl of SDS (1%) and HEPES buffer (20 nM, pH 7.9), boiled at 90°C for 5 minutes, vortexed briefly, and allowed to cool on ice for 3 minutes. Labeling buffer was then added followed by UDP-GalNAz and Gal-T1. After overnight incubation at 4°C, the GalNAz-labeled O-GlcNAc-modified protein mixture was precipitated using the chloroform/methanol precipitation method and resuspended in 50µl of buffer containing SDS (1%) and Tris (50mM, pH 8). The sample was subsequently labeled with the TAMRA-alkyne dye by using the Click-It™ TAMRA Glycoprotein Detection Kit (Thermo Fisher Scientific, Waltham, MA). The mixture was vortexed for 5 seconds after the addition of each component according to manufacturer’s instructions. The mixture was then rotated for 1 hour at 4°C for the conversion of the azide group to a stable triazole conjugate (TAMRA). DTT (25 mM) was added to stop the reaction. Labeled samples were precipitated using methanol/chloroform/water, brought up to a concentration of 2 mg/mL in 1% SDS and HEPES buffer (20 nM, pH 7.9), plus Complete™ protease inhibitors.

### Immunoprecipitation of TAMRA-Labeled O-GlcNAc Proteins

TAMRA labeled proteins were immunoprecipitated using Pierce MS-Compatible Magnetic IP Kit (Thermo Fisher Scientific, Waltham, MA) according to manufacturer’s instructions. Briefly, the labeled protein solution was precleared against washed protein A/G magnetic beads (20 µL/200 µg of protein) at 4 °C for 1 h. After bead separation, the supernatant was collected and incubated with an anti-TAMRA antibody (Thermo Fisher Scientific, Waltham, MA) at 10 μg/200 µg of protein at 4 °C overnight. The samples were then added to pre-washed protein A/G magnetic beads (25 µL) for 1 hour. The beads were collected and washed three times with IP-MS Wash Buffer. After washing, the beads were mixed with 100 µl elution buffer for 10 min. The supernatant was collected and dried for MS analysis.

### Mass spectrometry

From the same hearts, we prepared and ran TAMRA-enriched samples for assessing O-GlcNAc protein levels along with unenriched samples for a comparison to global protein levels. We hereafter refer to the TAMRA-enriched proteins as O-GlcNAcylated proteins. The samples were reduced with 5 mM dithiothreitol at 37 °C for 1 h, and alkylated with 10 mM iodoacetamide in the dark at 25 °C for 45 min. Then 100 mM Tris HCl was added. Samples were first digested with LysC (FUJIFILM Wako Chemicals, Richmond, VA) at enzyme to substrate ratio of 1:50 at 25 °C for 2h, followed by trypsin (Promega Corporation Madison, WI) digestion overnight (c.a. 14 h) at 25 °C with enzyme to substrate ratio of 1:50. Digested peptides with TAMRA enrichment were vialed at 0.1 µg/µL in 0.1% formic acid for liquid chromatography mass spectrometry (LCMS). A Waters NanoAcquity was operated in direct injection mode. Mobile phase A (MPA) was 0.1% formic acid in 97:3 water: acetonitrile, and mobile phase B (MPB) was 0.1% formic acid in 10:90 water: acetonitrile. LC column dimension was 20 cm × 75 µm i.d. with 1.9 µm ReproSil C18 packing prepared in house. 5 µL of peptides were loaded at 2% MPB for 30 min. Separation was performed by a gradient of 6% to 30% MPB over 85 min, followed by 10 min ramp to 60%. Flow rate was constant at 0.2 µL/min and column temperature was at 50 °C. Mass spectra were collected using a Thermo Fisher Scientific Orbitrap Lumos (Waltham, MA) with data-dependent acquisition. MS resolution was 60k for MS1 and 30k for MS2. Alternating HCD (charge state 2-10) and ETD (charge state 3-10) were used for MS2. Data were processed with PEAKS Studio (10.0 build 2190129) with label free quantitation. Dynamic modifications included protein N-terminal acetyl, methionine oxidation, carbamidomethylation, phosphorylation, O-HexNAc, O-HexNAc with TAMRA label. False discovery rate filter was 1% with decoy-fusion option in PEAKS Studio. The protein abundances (total areas) were log2 transformed and normalized to median in Perseus [19]. Missing values were imputed from normal distribution. Raw data are available in massive.ucsd.edu (Accession: MSV000088525. Reviewer Password: Heart3976) and ProteomeXchange [20] with accession PXD030218.

### Bioinformatics analysis

The gene-annotation enrichment analysis software, DAVID Bioinformatics Resources 6.8 [21, 22], was utilized to acquire molecular function (MF) and KEGG (Kyoto Encyclopedia of Genes and Genomes) pathway [23] information on the protein sequences demonstrating significantly higher O-GlcNAc levels during POH.

### Immunoprecipitation (IP) to confirm O-GlcNAc levels on proteins of interest

Two metabolic enzymes identified as having increased O-GlcNAc levels with POH were chosen for further evaluation namely pyruvate dehydrogenase (PDH) and carnitine O-palmitoyltransferase 1, muscle isoform (CPT1B). We used the PDH Rodent Immunocapture Kit (Abcam, Waltham, MA) to examine the subunites comprising this complex. Briefly, frozen heart tissues were lysed with 1X Extraction Buffer with a protease inhibitor cocktail and the OGA inhibitors PUGNAc 20 µmol/L (Toronto Research Chemical, North York, ON, Canada) and Thiamet-G 1 µmol/L (Adooq Bioscience, Irvine, CA). Solubilized homogenate was immunocaptured by mixing with PDH antibodies coupled agarose beads for 3 hours at 4 C in a tube rotator. The beads were collected by centrifugation for 2 minutes at 300×g and washed three times. The immunoprecipitated proteins were eluted with 40-μl SDS Elution Buffer. The samples were separated by SDS-PAGE and transferred to onto a polyvinylidene fluoride membrane. The membrane was subsequently stained for total protein with Pierce™ Reversible Protein Stain Kit (Thermo Fisher, Waltham, MA). The membranes were probed with Anti-O-Linked N-Acetylglucosamine antibody (anti-RL2, Novus Biologicals, Littleton, CO). The immunoblots were visualized with enhanced chemiluminescence using the chemiDoc-It™ imaging system (Analytik Jenna USA LLC). O-GlcNAc levels on the PDH proteins were normalized to protein staining in the same molecular weight region.

To assess changes in CPT1B O-GlcNAc levels, we performed the immunoprecipitation following the instructions detailed in the Pierce Co-Immunoprecipitation Kit (#26149, Thermo Fisher Scientific, Waltham, MA). Briefly, the tissue lysate was precleared against washed protein A/G magnetic beads (20 µL/200 µg of protein) at 4 °C for 1 h. After bead separation, the supernatant was collected and incubated with an anti-RL2 antibody (Novus Biologicals, Littleton, CO) at 10 μg/200 µg of protein at room temperature for 1 hour. The samples were then added to pre-washed protein A/G magnetic beads (25 µL) for 1 h. The beads were collected and washed three times with IP-MS Wash Buffer. After washing, the immunoprecipitated proteins were recovered by boiling in 50 µl 1X SDS buffer. The samples were separated by SDS-PAGE and transferred onto a polyvinylidene fluoride membrane. The membranes were probed with CPT1B antibody (Proteintech, Rosemont, IL). The immunoblots were visualized with enhanced chemiluminescence using the chemiDoc-It™ imaging system (Analytik Jenna USA LLC). We normalized the input CPT1B protein levels to protein staining in the molecular weight region of interest. The IP fraction was normalized to input protein concentration as determined by the BCA assay (Thermo Fisher Scientific, Waltham, MA).

### CPT1 Enzyme activity assay

CPT1 enzyme activity was determined through monitoring thiolytic cleavage of palmitoyl-CoA. The rate of the thiolytic cleavage was measured spectrophotometrically at 412 nm as described by Bieber et al. [24]. In brief, 20 mg of heart tissue were lysed and homogenized in ice-cold buffer containing 0.25 M sucrose, 1 mM EDTA, 0.1% ethanol, and protease inhibitors cocktail. Total protein was quantified by BCA assay (Thermo Fisher Scientific, Waltham, MA). The enzymatic activity was measured in 175 µl reaction buffer (116 mM Tris-HCl, 2.5 mM EDTA, 2 mM 5,5’-dithio-bis-(2-nitrobenzoic acid) (DTNB), pH 8.0) and 10 µg lysates in a final volume of 188 µl by adding homogenization buffer. The samples were then incubated at 30°C for 5 min to eliminate reactive thiol groups and the resulting background absorbance was measured. After preincubation, 10 µl palmitoyl-CoA (2 mM prepared in double distilled water) and 2 µl L-carnitine solution (1.2 mM in 1 M Tris-HCl, pH 8.0) were added to the reaction mixtures. Kinetic reads were collected at 2 minutes intervals for 30 min. In a parallel experiment, 0.2 % Triton X-100 was added to inactivate CPT1 [2]. The CPT1 activity was calculated by subtracting the activity with Triton X-100 from total activity. Activity was defined as nmol CoA-SH released/min/mg protein. Calculations were made assuming a molar extinction coefficient of 13,600 M^−1^ cm^−1^ at 412 nm for the thiol reagent DTNB.

### PDH Enzyme activity assay

PDH enzyme activity was measured according to the manufacturer’s instructions for the enzyme activity assay kit (K679-100, BioVision Inc., Milpitas, CA). Briefly, heart lysate was prepared with PDH assay buffer plus protease inhibitors (78440, Thermo Fisher, Waltham. MA). 20 µg/50 µL of lysate was loaded onto microplate wells. After adding 50 µL PDH reaction solution, PDH activity was determined by monitoring the change in absorbance at 450 nm over 60 min at 37 °C. PDH activity was calculated by determining the slopes of the generated curves between the 10-min and 40-min time points where the increase in absorbance was linear. PDH Activity was defined as nmol NADH released/min/mg protein.

### Data Visualization and Statistical Analysis

We preformed data visualization using GraphPad Prism version 8.3.3 (GraphPad Software, San Diego, CA, USA). All reported values are means ± standard error of the mean (SEM). All experiments except the TAMRA-enriched (O-GlcNAc) proteomics used an unpaired two-tailed T-test. We initially used a two-tailed T-test to analyze the TAMRA-enriched (O-GlcNAc) samples and found that all significant changes were increased for POH over Sham, so we were justified to use a one-tailed T-test for this experiment only. Criterion for significance was p < 0.05.

## RESULTS

### Echocardiographic and morphometric results during POH

We performed echocardiograms to show the cardiac parameters and the degree of aortic constriction (Figure 1). The aortic constriction led to an average peak instantaneous pressure gradient of 79.6±7.7 mmHg across the constriction site for POH. Left ventricular chamber size and systolic function was similar between the groups. As expected, heart weight to tibial length (HW/TL) was significantly elevated in POH confirming the presence of left ventricular hypertrophy (Figure 1).

**Figure 1.**
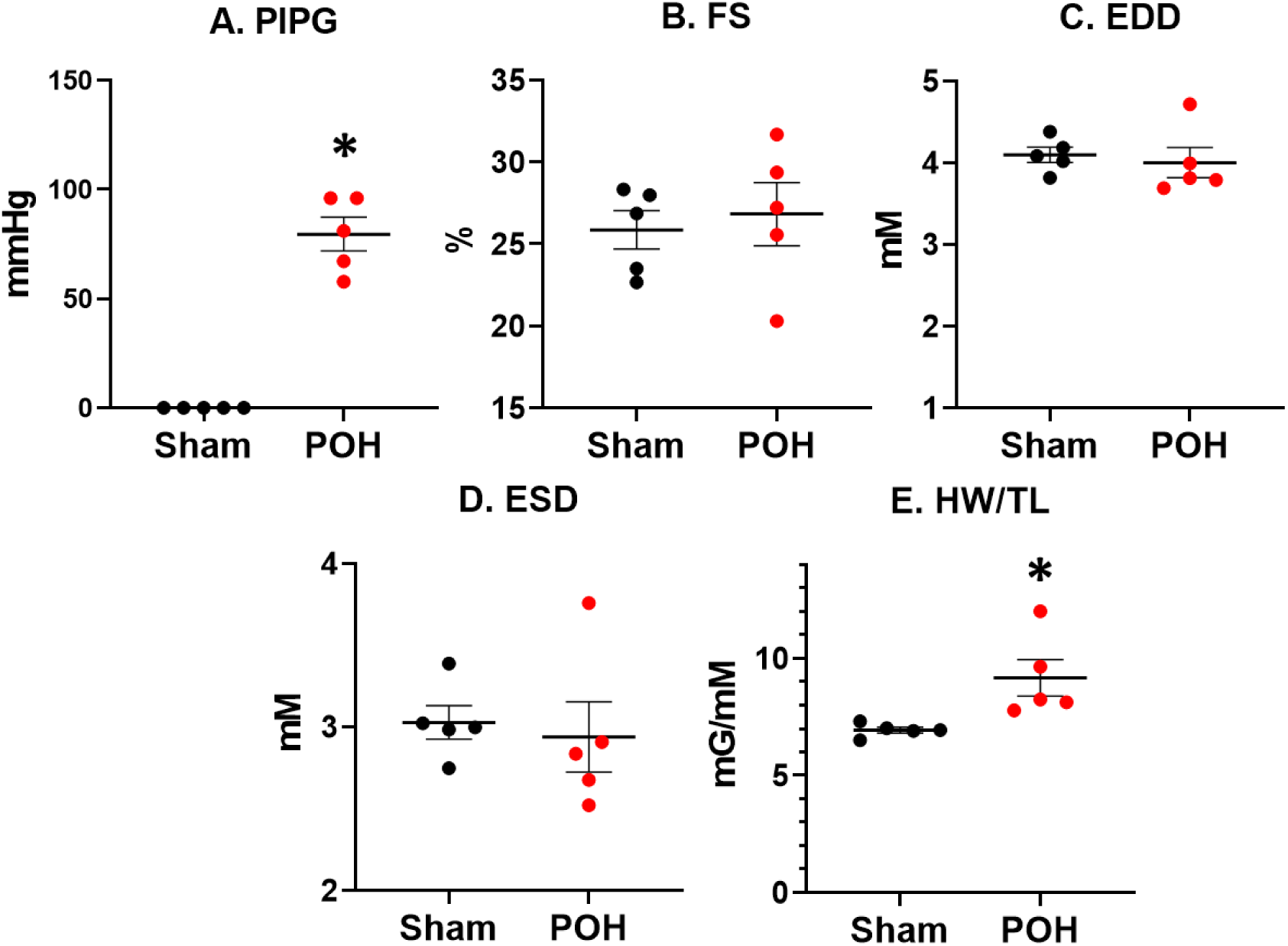
Echocardiogram and morphometrics in pressure-overload hypertrophy (POH) and sham (Sham) groups. Bars indicate mean± standard error of the mean. * designates p<0.05 between the indicated groups. PIPG, peak instantaneous pressure gradient across the transverse aortic constriction site; FS, left ventricular fractional shortening; EDD, left ventricular end-diastolic diameter; ESD, left ventricular end-systolic diameter; HW/TL, heart weight normalized to tibial length; mmHg, millimeter mercury; mM, millimeter; mG, milligram. N=5 for all groups.

### Protein Identification with analysis of biological processes and KEGG pathways

We identified a total of 707 O-GlcNAcylated proteins in Sham and POH (Supplemental Table 1). Of these, 233 O-GlcNAcylated proteins were significantly increased in POH over Sham whereas no proteins were significantly decreased in POH (Supplemental Table 1 and 2). We found 26 proteins that were significantly increased for both O-GlcNAc and global protein levels, however the fold change was higher for O-GlcNAc in all but one of these proteins (Supplemental Table 3).

We used the DAVID software to analyze the proteins with significantly increased O-GlcNAc levels to gain insight into the biological processes and signaling pathways altered. A cutoff Benjamini corrected p-value of less than 0.05 was used in Tables 1 and 2. The biological processes and KEGG pathway analysis identified proteins predominantly involved in metabolic and structural roles. Supplemental Tables 4 and 5 include the list of proteins with increased O-GlcNAc levels within each category.

**Table 1.**
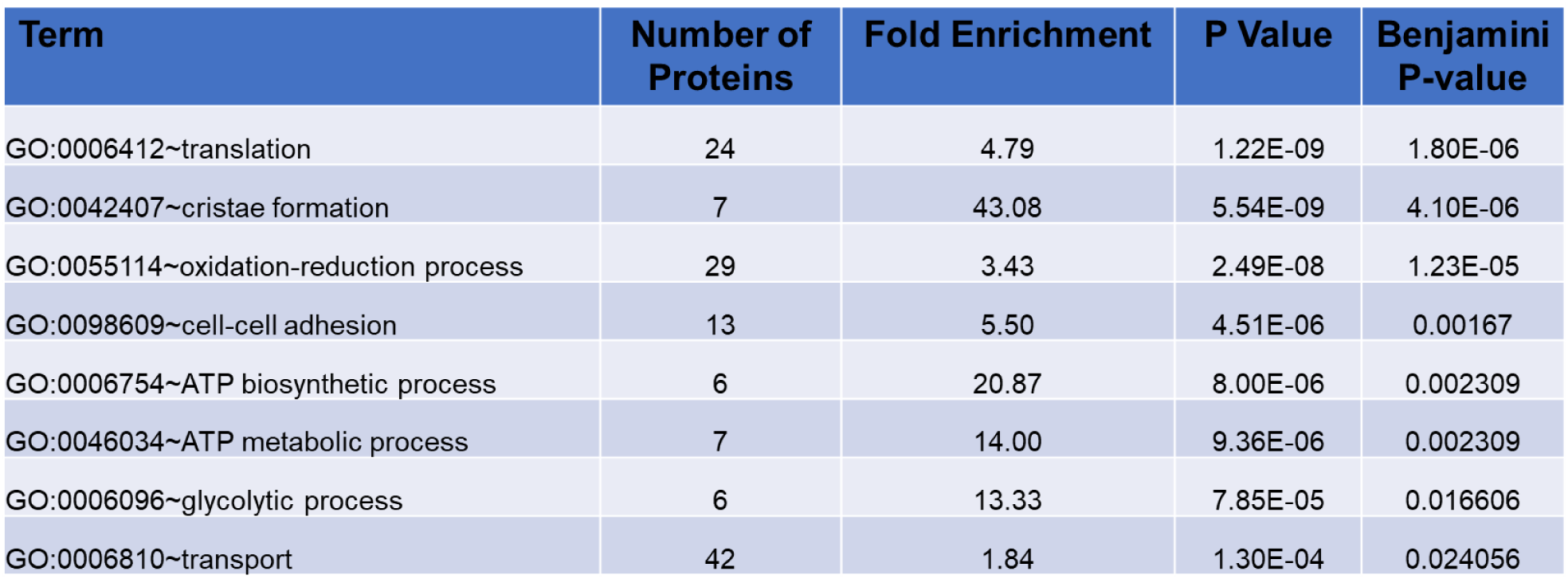
Analysis of proteins with significantly increased O-GIcNAc levels during pressure overload hypertrophy (POH) for biological processes.

**Table 2.** Analysis of proteins with significantly increased O-GIcNAc levels during pressure overload hypertrophy (POH) for KEGG pathways.

| Term | Number of proteins | Fold Enrichment | P Value | Benjamini P-value |
| --- | --- | --- | --- | --- |
| mmu01200:Carbon metabolism | 17 | 7.88 | 2.81E-10 | 4.76E-08 |
| mmu00190:Oxidative phosphorylation | 18 | 6.96 | 5.18E-10 | 4.76E-08 |
| mmu01130:Biosynthesis of antibiotics | 20 | 5.03 | 1.12E-08 | 5.42E-07 |
| mmu05012:Parkinson's disease | 17 | 6.14 | 1.18E-08 | 5.42E-07 |
| mmu05016:Huntington's disease | 19 | 5.16 | 1.94E-08 | 7.13E-07 |
| mmu03010:Ribosome | 16 | 5.93 | 5.76E-08 | 1.77E-06 |
| mmu05010:Alzheimer's disease | 17 | 5.17 | 1.37E-07 | 3.59E-06 |
| mmu00010:Glycolysis / Gluconeogenesis | 11 | 8.96 | 2.93E-07 | 6.73E-06 |
| mmu01100:Metabolic pathways | 47 | 1.99 | 1.96E-06 | 4.01E-05 |
| mmu04932:Non-alcoholic fatty liver disease (NAFLD) | 14 | 4.80 | 5.83E-06 | 1.07E-04 |
| mmu00620:Pyruvate metabolism | 7 | 9.65 | 7.04E-05 | 0.001177 |
| mmu00280:Valine, leucine and isoleucine degradation | 7 | 6.85 | 4.91E-04 | 0.007526 |
| mmu00630:Glyoxylate and dicarboxylate metabolism | 5 | 9.27 | 0.001848 | 0.026082 |
| mmu00071:Fatty acid degradation | 6 | 6.59 | 0.001984 | 0.026082 |
| mmu00020:Citrate cycle (TCA cycle) | 5 | 8.40 | 0.00268 | 0.03288 |
| mmu04260:Cardiac muscle contraction | 7 | 4.89 | 0.002882 | 0.033143 |

### Immunoblotting and enzyme activities

We chose two metabolic enzymes with increased O-GlcNAc levels from the proteomics database to validate O-GlcNAc status via a separate method along with an assessment on how the change in O-GlcNAc levels affects enzyme activity. CPT1B was chosen due to its function as a shuttle of long chain fatty acids into the mitochondria for beta-oxidation. First, CPT1B protein expression in tissue lysate, which we denote as the input levels for IP, was higher in Sham versus POH (Figure 2). Post-immunoprecipitation using the RL2 antibody to capture O-GlcNAcylated proteins showed the CPT1B levels were greater in POH versus Sham confirming higher CPT1B O-GclNAcylation despite lower total levels during POH. We decided against using these experimental groups to evaluate CPT1 enzyme activity because the differences in total CPT1B would confound the results. Thus, we evaluated CPT1B in V-TAC and TG-TAC hearts, which we previously demonstrated have significantly different global O-GlcNAc levels (higher in TG-TAC). Total CPT1B levels were similar in V-TAC versus TG-TAC (denoted as the input for the IP), however CPT1B was greater in TG-TAC after IP indicating higher CPT1B O-GlcNAcylation for TG-TAC. We found CPT1 enzyme activity was higher in TG-TAC versus V-TAC demonstrating O-GlcNAcylation increases CPT1 activity (Figure 3).

**Figure 2.**
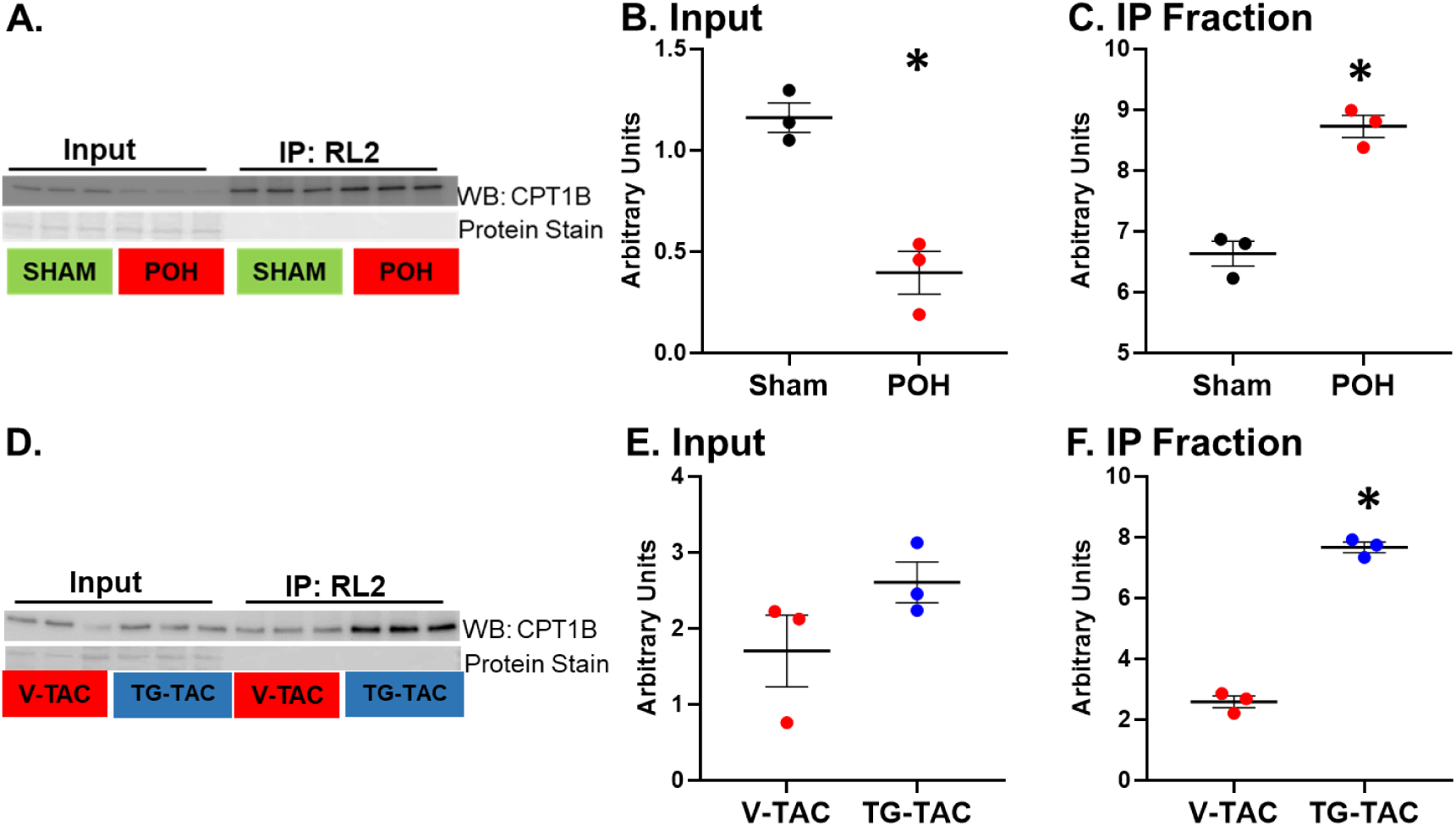
O-GlcNAc immunoprecipitation (IP) for carnitine palmitoyltransferase 1B (CPT1B). CPT1B western blots are show prior to (input) and after IP with the anti-O-GlcNAc antibody RL2 in POH and Sham (A) with quantification for input (B) and from IP (C). Western blots for experimental groups V-TAC and TG-TAC are shown in D with quantification for input (E) and from IP (F). Bars indicate mean± standard error of the mean. * indicates p<0.05 between the indicated groups. N=3 per group.

**Figure 3.**
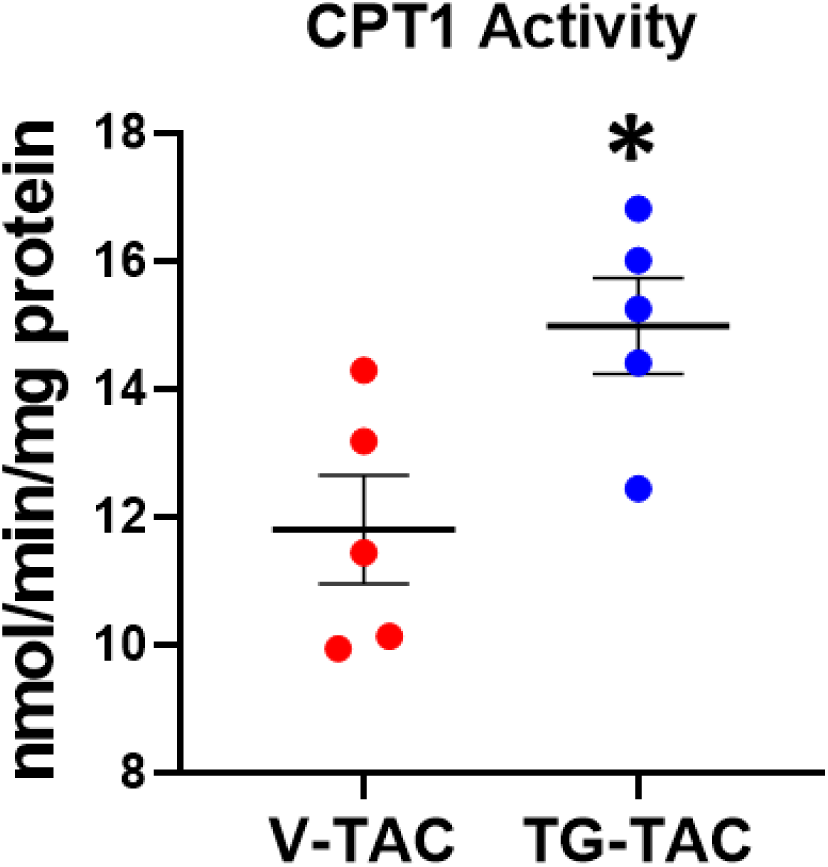
Quantification of enzyme activity for CPT1. Activity is defined as nmol CoA-SH released/min/mg protein. Bars indicate mean± standard error of the mean. indicates p<0.05 between the indicated groups. N=5 per group.

Next, we evaluated the PDH complex. Our proteomics results showed increased O-GlcNAcylation of the PDH components dihydrolipoyl dehydrogenase and dihydrolipoyllysine-residue acetyltrasferase. We immunoprecipitated the PDH complex and then performed a RL2 western blot on the bound fraction to determine O-GlcNAc levels. We found increased O-GlcNAc levels on the immunoprecipitated PDH complexes in POH compared to Sham (Figure 4). Utilizing the V-TAC versus TG-TAC model, we assessed PDH enzyme activity and found decreased PDH enzyme activity in TG-TAC hearts suggesting that higher O-GlcNAcylation inhibits PDH activity (Figure 5).

**Figure 4.**
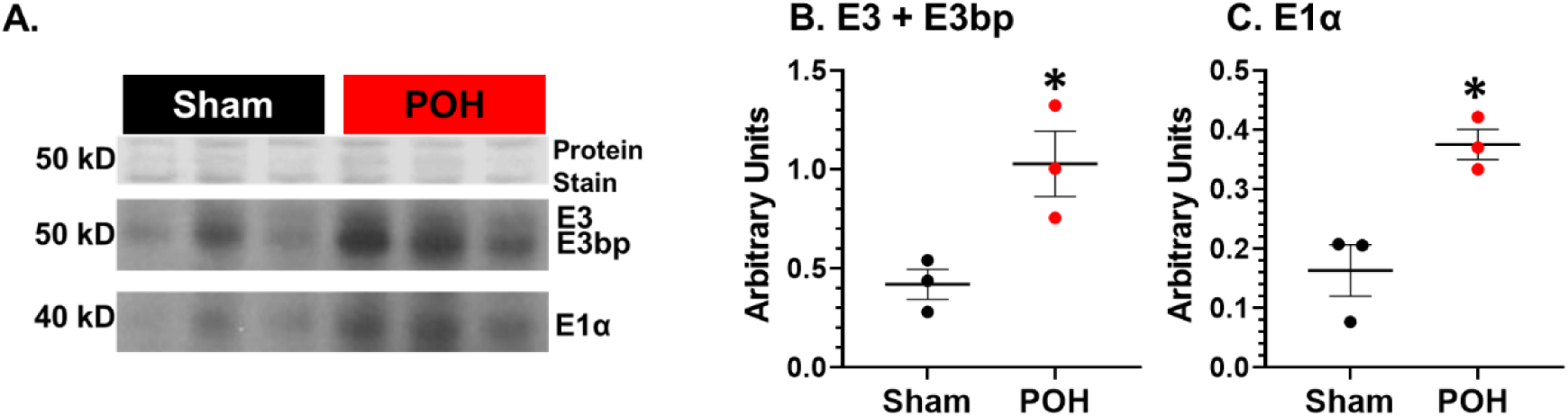
O-GlcNA levels on pyruvate dehydrogenase (PDH) immunocaptured proteins. PDH complex immunocaptured proteins underwent western blot with the anti-O-GlcNAc antibody RL2. (A) shows protein staining for the immunocaptured proteins and the RL2 bands at 50 kD and 40 kD. The bands at 50 kD correspond to the subunits E3 and E3 binding partner (E3bp) whereas 40 kD corresponds to the E1α subunit. (B) Quantification of E3 and E3bp normalized to protein staining. (C) Quantification of the E1α subunit normalized to protein staining. Bars indicate mean± standard error of the mean. * indicates p<0.05 between the indicated groups. N=3 per group.

**Figure 5.**
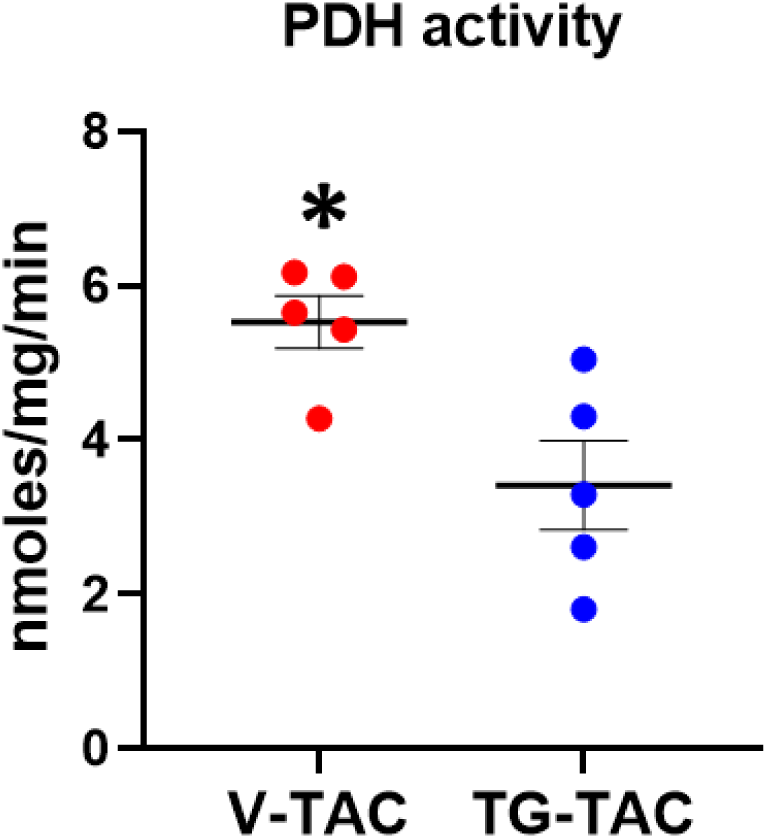
Quantification of enzyme activity PDH. Bars indicate mean± standard error of the mean. * indicates p<0.05 between the indicated groups. N=5 per group.

## DISCUSSION

We previously knew only a few O-GlcNAc modified proteins and functional effects during POH primarily because of their overall low abundance and because the fragile O-GlcNAc moiety is lost during standard LC-MS analysis used for large-scale protein identification. Utilizing Click-iT™ technology, we bypassed this technical problem to identify a total of 707 O-GlcNAcylated proteins in Sham and POH, 233 of which exhibited significantly increased O-GlcNAcylation during POH. No proteins had significantly decreased O-GlcNAc levels during POH. Thus, we have generated the first comprehensive list of proteins with changes in O-GlcNAc levels during POH and, to our knowledge, the largest list for any cardiac pathology. Our results demonstrate the large number of proteins and cellular processes affected by O-GlcNAc during POH and serve as a guide for testing specific O-GlcNAc-regulated mechanisms.

O-GlcNAcylation is often described as beneficial for acute stressors and detrimental with prolonged elevations [25]. However, because of the large number of affected proteins, it is conceivable O-GlcNAc changes during POH are beneficial for some proteins/cellular functions while detrimental for others. Accordingly, we recommend future studies evaluate O-GlcNAc’s effects on specific proteins and cellular functions during POH. This experimental approach could potentially optimize future translational therapies by identifying specific O-GlcNAcylated protein to target rather than global levels. Emerging technologies like aptamers or nanobodies could be developed to modify O-GlcNAc levels at specific protein sites [1]. It may also be possible to modify select groups of O-GlcNAcylated proteins by altering activity of the accessary proteins that target OGT to specific proteins [1].

### O-GlcNAcylated proteins and cellular functions during POH

Previous findings from mostly non-cardiac tissues indicates protein O-GlcNAcylation affects many cellular functions including cell cycle regulation, transcriptional and translational events, metabolism, mitochondria function, protein synthesis and quality control, autophagy, epigenetics, cellular signaling, contractile and cytoskeletal proteins, and calcium-handling [26, 27]. In our study, proteins with increased O-GlcNAcylation were overrepresented for similar biological processes like translation, metabolism (ATP biosynthesis and metabolic processes, glycolytic processes), mitochondrial function (cristae formation) and cytoskeletal proteins (cell-cell adhesion). KEGG pathway analysis showed overrepresentation for multiple metabolic pathways, ribosome function and cardiac muscle contraction. During POH, ribosome function could affect hypertrophic growth and proteostasis whereas metabolic pathways and cardiac muscle contraction could directly alter cardiac function.

It is also noteworthy that many metabolic pathways including glycolysis, glucose/pyruvate oxidation, and fatty acid oxidation (FOA) were highly enriched for proteins with increased O-GlcNAcylation because this PTM is frequently described as a nutrient sensor linking metabolic changes to protein function [2, 6]. The hexosamine biosynthesis pathway (HBP) branches from glycolysis to produce UDP-GlcNAc, the moiety used by OGT for O-GlcNAc PTMs. OGT is sensitive to UDP-GlcNAc concentration making overall O-GlcNAc levels responsive to changes in HBP flux [15]. Thus, it is widely assumed that greater glycolytic rates, as occurs in POH or heart failure, increase HBP flux to augment overall protein O-GlcNAc levels [6, 16]. However, we recently measured HBP flux in the heart for the first time and found that HBP flux does not change even through a 20-fold variation in glycolytic rates [28]. Our findings herein expand upon our prior work and suggest HBP flux and O-GlcNAc levels alter downstream metabolic enzymes which determine cardiac fuel metabolism. Whereas the unstressed heart relies mainly on fatty acid oxidation (FAO) as a fuel source, POH augments glucose utilization [29, 30]. This metabolic shift is important as changes in glucose metabolism directly affect hypertrophic growth and augmenting FAO in POH may inhibit progressive cardiac dysfunction [31–33]. Yet, strategies targeting fuel utilization for the CAC to prevent heart failure in POH have not been realized partially because the mechanisms underlying these metabolic changes are incompletely known. Thus, it is important the determine the contribution of O-GlcNAc to the metabolic alterations occurring during POH.

In this regard, we assessed two metabolic enzymes, PDH and CPT1B, key enzymes for glucose/pyruvate oxidation and fatty acid metabolism respectively. We used immunoprecipitation to confirm the LC-MS results showing increased O-GlcNAcylation for both enzymes during POH. Although O-GlcNAcylation increased, CPT1B total protein levels were lower in POH versus Sham thus confounding assessment of enzyme activity between these groups. Therefore, we used the TG-TAC and V-TAC hearts for both enzyme activity assays. Higher O-GlcNAcylation decreased PDH activity whereas it augmented CPT1B activity. O-GlcNAc’s effect on PDH could contribute to the finding that glucose oxidation often does not increase to the same extent as glycolysis during POH [29]. We speculate CPT1B O-GlcNAcylation could help preserve FAO in the face of alterations in other FAO enzymes occurring during POH [30]. Of course, O-GlcNAcylation increased on multiple enzymes involved in glucose and fatty acid metabolism; the citric acid cycle; and even mitochondrial function, so a more detailed metabolic characterization is necessary.

It is notable but expected that not all O-GlcNAcylated proteins had increased levels with POH. We do not know the mechanisms regulating O-GlcNAcylation on specific proteins during POH. OGT, the single protein responsible for attaching the O-GlcNAc moiety to proteins, has binding partners and/or PTMs that may confer specificity [1]. Future studies should evaluate how these factors impact OGT protein targeting during POH. It would also be important to know whether the existing transgenic mice used in studies to alter myocardial O-GlcNAc levels during POH broadly affect all O-GlcNAcylated proteins or just the subset of proteins changing during POH because these differences could have profound effects on cardiac outcomes.

### Limitations

Our approach relies on protein level enrichment and identification of primarily non-modified peptides derived from these enriched proteins using LC-MS. Hence, it is possible that some of the proteins identified with our TAMRA enrichment protocol are not O-GlcNAcylated, rather they bind to the O-GlcNAcylated proteins during the TAMRA IP or represent other protein interferences. Indeed, this is a common challenge when enriching low abundant proteins from a complex mixture using affinity purification methods. Thus, future studies should use another method to confirm O-GlcNAcylation status of our reported proteins. Furthermore, our methodology does not resolve site specificity for the O-GlcNAc modification. Site specific data would provide additional proof that an identified protein is O-GlcNAcylated along with determining whether an individual protein is modified on more than one site. Site specific information is also necessary for using gene editing techniques to remove O-GlcNAc sites while studying a protein of interest.

### Summary

We generated the first comprehensive list of proteins with changes in O-GlcNAc levels during POH. Our list serves as a guide for testing specific O-GlcNAc-related mechanisms affecting the heart during POH.

## ACKNOWLEDGEMENTS

We thank Ronald Moore and Matthew Monroe for helping with proteomics experiments and proteomics data deposition, respectively. A portion of this research was performedcat the Environmental Molecular Sciences Laboratory, a DOE Office of Science User Facility sponsored by the Biological and Environmental Research program under Contract No. DE-AC05-76RL01830.

## GRANTS

Research reported in this publication was supported by the National Heart, Lung, and Blood Institute of the National Institutes of Health under award number NIH R01HL122546 to AKO and NIH K01HL120886 to DRL.

## Notes

ftp:///

http://proteomecentral.proteomexchange.org/cgi/GetDataset?ID=PXD030218

### Competing Interest Statement

The authors have declared no competing interest.

